# Reduced ribosome activity influences the non-uniform evolution of 16S rRNA hypervariable regions

**DOI:** 10.1101/2022.09.23.509299

**Authors:** Nikhil Bose, Sean D. Moore

## Abstract

16S rRNA gene sequences are commonly analyzed for taxonomic and phylogenetic studies because they contain hypervariable regions that can help distinguish different genera. However, intra-genus distinction is often difficult due to high sequence identities among closely related species. Although common tools for 16S sequence taxonomic classification weight residue variations equally during comparisons, specific residues within hypervariable regions have not drifted evenly through evolution, suggesting that portions of them may be biologically important. We developed an *in vivo* test system where 16S variants coexisted among natural ribosome populations which allowed their fitness to be evaluated. We found that versions with evolutionarily disparate hypervariable regions were underpopulated in ribosomes and active translation pools, even for a single nucleotide polymorphism (SNP), which indicates functional constraints to the free evolutionary drift of hypervariable regions. Using an *in silico* method (positional relative entropy), we analyzed over 12,000 16S V3-V4 sequences across *Escherichia* and *Shigella* strains and identified species that can be distinguished by position-specific SNPs present in multiple 16S alleles in a genome. When we evaluated these informative SNPs with our *in vivo* system, we discovered that ribosomes harboring them were compromised, suggesting that their evolution is indeed biologically constrained. Overall, this study demonstrates that SNPs within hypervariable regions are not necessarily inconsequential and that common computational approaches for taxonomic 16S rRNA sequence classification should not assume an even probability of residues at each position.

**Importance:** Hypervariable regions within 16S rRNA genes are commonly analyzed to determine microbial diversity. However, because sequences within a genus are highly similar, strain- or species-specific identification is often uncertain. Because there are no established functions of hypervariable regions, residue variations within them are often evenly weighted when making taxonomic comparisons. This study established that 16S rRNAs with naturally occurring variations in hypervariable regions can affect ribosome quality, indicating that their residues should not be weighted equally during taxonomic sequence classifications.

## Introduction

16S ribosomal RNAs (rRNA) are found in all bacteria and archaea because they are required for small subunit assembly, ribosome formation (1–5), and for ribosomes to properly decode mRNAs (6, 7). The *rrs* genes that encode 16S rRNAs have diverged over time, with biochemically critical residues changing infrequently and hypervariable regions exhibiting greater sequence diversity (8, 9). Shannon entropy has been used to computationally evaluate the conservation of 16S gene residues across bacteria (10, 11), which confirmed the presence of nine hypervariable regions (V1-V9). These hypervariable region sequences are commonly analyzed for microbial classification, with the combined V3-V4 segment being the most common because its length accommodates affordable second-generation sequencing technologies (12–16). However, categorizing 16S sequences for a genus into specific species is not regularly achieved because the few (but potentially informative) residue differences among sequences are computationally outweighed by their overwhelming similarities (11, 17). This problem is exemplified when 16S gene analyses co-classify sequences as *Escherichia* and *Shigella* (18–20), even though residue differences among species exist (21, 22). Applying stringent sequence classification criteria (23, 24) and incorporating more 16S rRNA sequences into databases can partially alleviate this problem, but a lack of consideration for intra-genome allelic variation makes such an endeavor less effective. Because relatedness is commonly assigned by overall sequence similarity, the potential for individual residue identities across multiple 16S gene alleles to indicate considerably larger evolutionary divergence is overlooked. Moreover, the biological impact of 16S gene allelic variations in a genome is often ignored.

Computational tools that help classify RNA sequences based on covariance models showed that co-variation of rRNA residues is important for secondary structures (25–28), hence residue deviations, even in hypervariable regions, may be biologically important. Orthogonal expression studies investigating *E. coli* 16S rRNA residue changes showed that certain conserved region SNPs can affect ribosome function (29, 30). One study explored the effects of random 16S rRNA mutations and identified a few variable region mutations as “mildly influential” to *E. coli* growth (31). Though the effects of these mutants on ribosome activity were not explored, that study suggests that variable region mutation can have a biological influence. However, the introduced mutations were not species-specific, and those bacteria were already stressed by a high growth temperature and high 16S rRNA expression levels. A recent study exploring reporter protein expression by orthogonal 16S rRNA variants showed varying translation activities for *Shigella* 16S sequences (32), which demonstrated that not all *Escherichia* and *Shigella* 16S sequences have equal functional capabilities. However, the impact of individual variable regions and SNPs within them were not tested.

In this study, we evaluated 16S rRNA mutants that were expressed at a low level in *E. coli* containing unaltered chromosomal operons. Our system involved measuring the abundance of a plasmid-expressed tagged 16S rRNA mutants relative to chromosome-derived 16S rRNAs in ribosomes at various stages of assembly and translation. 16S rRNAs harboring SNPs at functionally conserved decoding center residues were less abundant in polysomes, which is consistent with their established biological importance. Surprisingly, we discovered that an *E. coli* 16S rRNA harboring a variable region 3 (V3) SNP exhibited relative abundances as low as that observed for a truncated V3 sequence from *Clostridioides difficile*. Using computational analyses, we determined the relative entropy of each residue position across over 12,000 V3-V4 variable region sequences from strains within closely related genera *Escherichia* and *Shigella* and identified polymorphisms that were prevalent across strains in certain species, indicating a greater evolutionary divergence for those species than their overall rRNA homology suggests. The *in vivo* performance of ribosomes harboring informative V3-V4 region polymorphisms was variable, including variations that distinguish *Escherichia albertii, Shigella dysenteriae* and *Shigella boydii*. Taken together, our computational and biological evaluations of 16S variable region polymorphisms support the model that biological functionality is tailoring their evolutionary drift and that simply comparing sequence identities overlooks important taxonomic information.

## Results

### 1. Development of a tagged V1 16S rRNA

We developed a 16S expression plasmid and evaluated variants in a *ΔrecA Escherichia coli* strain with unaltered chromosomal 16S genes. We removed *recA* to avoid recombination between plasmid and chromosomal *rrs* alleles. The plasmid included an *E. coli* 16S gene *rrsA* with a unique variable region 1 (V1) sequence that could be distinguished from chromosomal 16S genes by PCR specificity (**Fig. 1A**). A previous study with an engineered *E. coli* lacking chromosomal *rrn* operons demonstrated that a plasmid expressing 16S rRNA with a modified V1 region supported growth (33). Our sequence modification was designed such that the secondary structure of the V1 region in the expressed 16S rRNA would be similar to that for WT 16S rRNA (**Fig. S1A**). We confirmed that our primers detected the tagged V1 and WT V1 sequences separately (**Fig. S1B**). Mutations and variations were introduced to this parental tagged V1 *rrsA* for functional evaluation.

**FIG 1:**
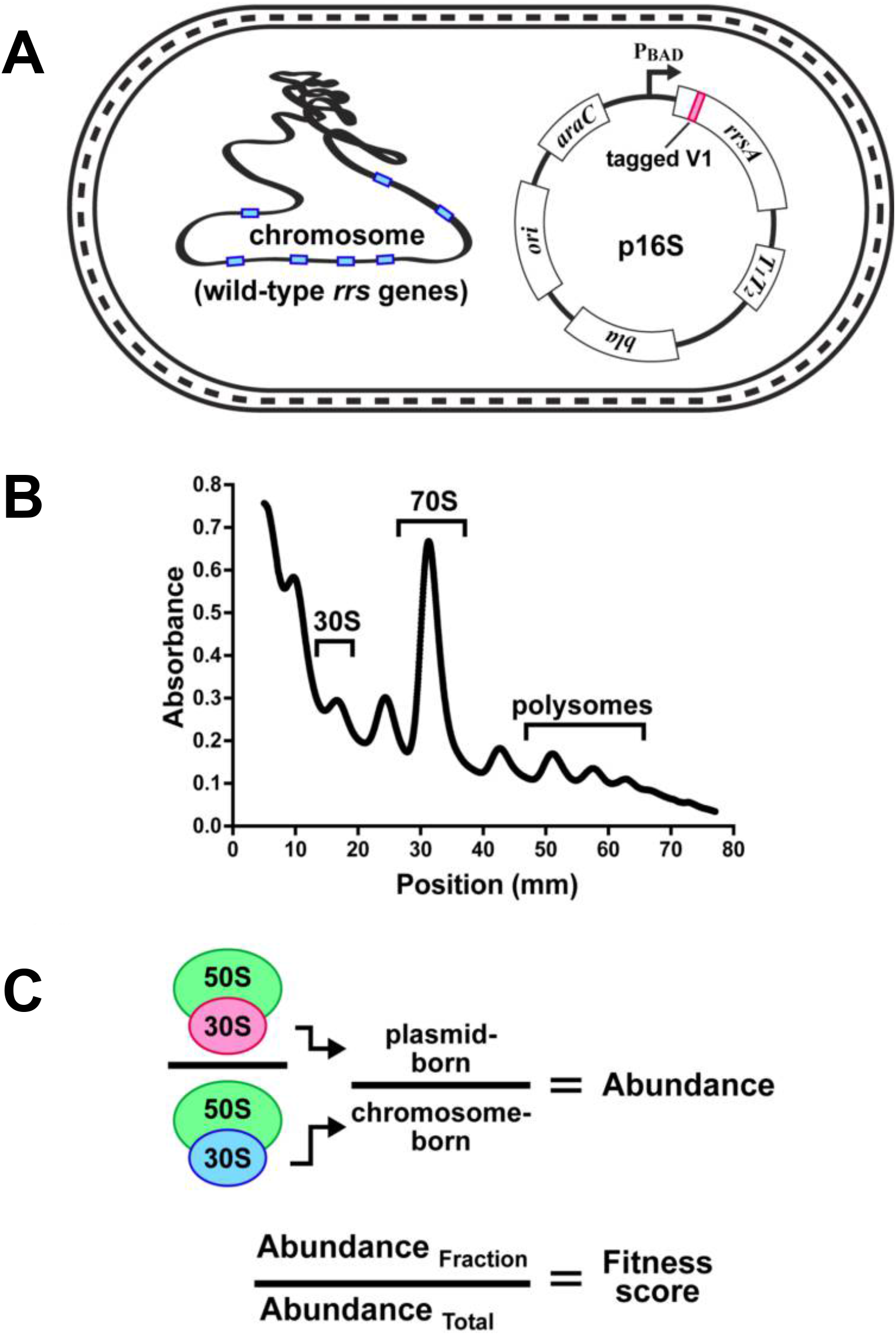
Establishing the fitness of 16S rRNA variants. Modified 16S rRNAs were evaluated in a wild-type *E. coli strain* with intact *rrn* operons. **A)** The *E. coli rrsA* gene was cloned into a plasmid under the control of a tightly repressed P_BAD_ promoter. The cloned *rrsA* was modified in its variable 1 (V1) region to contain a unique tracking tag sequence that was detectable using RT-qPCR. Other mutations were subsequently introduced in this tagged V1 *rrsA* for fitness evaluations. **B)** Fractionation of cell lysates using sucrose gradients allowed isolation of 16S rRNAs in various stages of assembly and translation. The regions of 30S, 70S, and polysome material collected in this study are indicated. **C)** RNA was extracted from gradient fractions and used to establish the abundance of plasmid-born 16S relative to chromosome-born in the same fraction. A fitness score was then calculated by comparing the abundance in a given fraction to that present in the total lysate.

Our experimental strategy involved growing strains harboring 16S variants in plasmids in a growth medium promoting low expression of the plasmid-born 16S rRNA relative to chromosome-born 16S rRNA. 16S rRNAs in immature or recycling small subunits were recovered from the 30S gradient position, mature RNAs present in monosomes were recovered from the 70S position, and RNAs participating in translation were recovered from polysomes having three or more ribosomes (**Fig. 1B**). The relative abundances of plasmid-born *vs*. chromosome-born 16S rRNAs in those fractions and also the total lysates were determined using RT-qPCR. The persistence of the expressed variant 16S rRNA in ribosomes was determined by evaluating a “fitness score” which is the ratio of the abundance in a gradient fraction relative to the abundance in the total lysate. Fitness scores were then compared to scores obtained for the parental 16S with a tagged V1 lacking any other sequence changes (**Fig. 1C**).

### 2. Detection of plasmid born 16S rRNAs in ribosomes

We evaluated the expression and performance of plasmid-born, tagged parent 16S rRNA by determining its abundance and fitness scores after it had been expressed at a very low level (repressed by supplemented glucose), at somewhat higher levels (glycerol supplemented), and at a very high level (arabinose supplemented). The plasmid-born 16S rRNA was detected in 30S, 70S, and polysome fractions for all expression tests. The overall expression levels followed expectation, with the total abundance of the plasmid-born relative to the chromosomally-born version ranging from approximately 1:2000 (repressed), to approximately 1:400 (uninduced), to approximately 1:3 (induced) (**Fig. 2A**). Sanger sequencing of the qPCR products confirmed presence of the tagged V1 in those amplicons.

**FIG 2:**
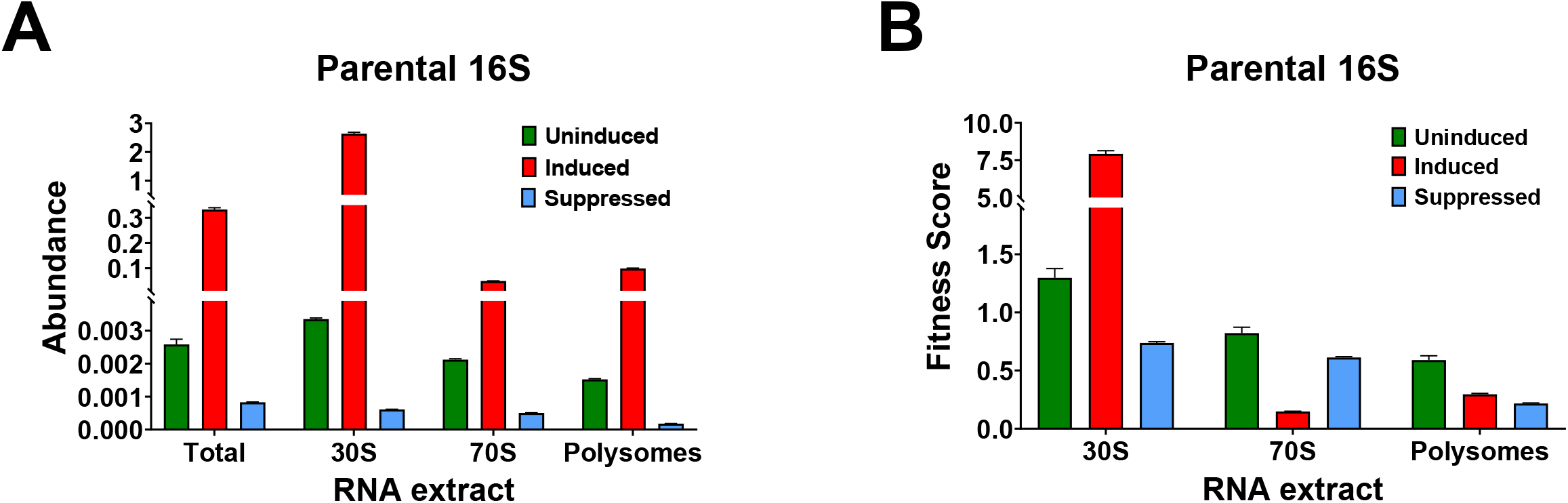
Performance assessment of parent 16S for various expression conditions. The plasmid-born 16S rRNA was detected by RT-qPCR in 30S, 70S, and polysome pools for uninduced, induced, and repressed culture growth. **A)** The abundance of plasmid vs. chromosome-born 16S RNA was determined for RNA extracts from total cell lysates (Total), ribosome small subunit assembly (30S), ribosome formation (70S), and translating ribosomes (polysomes). **B)** The abundances in each ribosome fraction were divided by their respective total cell lysate abundances to obtain fitness scores. Error bars represent standard deviation for biological replicates (n=3).

Fitness scores for the 16S rRNAs synthesized at high levels were calculated to be ~7.9 for 30S material (indicating a strong over-representation), ~0.15 for 70S, and ~0.29 for polysomes (indicating strong under-representations) (**Fig. 2B**). The high 30S scores suggest that a large proportion of the plasmid-expressed 16S rRNA was trapped in small subunit assembly, likely because the over-expression had depleted assembly or maturation resources. In support of this conclusion, overexpressing this 16S variant substantially impeded culture growth rate and it was significantly more toxic than the overexpression of a non-coding RNA of similar length (**Fig. S2**). The low fitness scores for the 70S and polysome fractions likely resulted from an overall impediment to ribosome production upon overexpression, such that pre-existing, untagged particles remained dominant in the translation pool. In comparison, cultures with uninduced expression exhibited faster growth rates and higher 16S rRNA fitness scores ranging from ~1.29 for 30S to ~0.6 for polysomes. Suppressed expression scores ranged from ~0.73 for 30S to ~0.22 for polysomes. Here, disparities in the fitness scores across the fractions may have been caused by some influence of the altered V1 region or from variable expression levels during culturing such that more tagged particles were produced later in the experiment. Therefore, it became important for mutant evaluations to be performed using the same culturing times and harvest windows as the parental reference cultures. The highest fitness scores in 70S and polysomes were achieved for the uninduced condition, making it the preferred expression condition to evaluate variant 16S rRNAs. These results showed that the RT-qPCR strategy could detect low-levels of tagged 16S rRNAs and that the plasmid-born version was incorporated into functional ribosomes. Also, having the ability to detect very low levels of 16S rRNA provided an opportunity to evaluate polymorphisms that might compromise ribosome formation and bulk translation if expressed at higher levels.

### 3. Performance evaluation of decoding center mutants

We sought to establish references for low fitness scores by expressing 16S rRNA variants harboring defects in critical areas. A prior report indicated that small subunits with decoding residue mutations A1492U, A1493U, or G530C were incapable of synthesizing a reporter protein (29); the transcript presence in mature ribosomes was not detectable, suggesting that that over-expression system may have additionally compromised maturation. Our system allowed us to assess abundances of decoding residue mutants without interfering with the function of abundant wild-type ribosomes. These decoding center mutations were introduced independently into the tagged plasmid 16S gene and expressed under an uninduced condition to obtain fitness scores. Each mutant 16S rRNA was detected in the 30S, 70S, and polysome pools. Fitness scores for 30S and 70S were higher relative to the parent 16S for A1492U and A1493U, while polysome scores for all mutants were significantly lower, especially for A1493U and G530C (**Fig. 3**). These results demonstrated that 16S rRNAs with these decoding center mutations could form mature particles and enter polysome pools, although their presence there was notably reduced.

**FIG 3:**
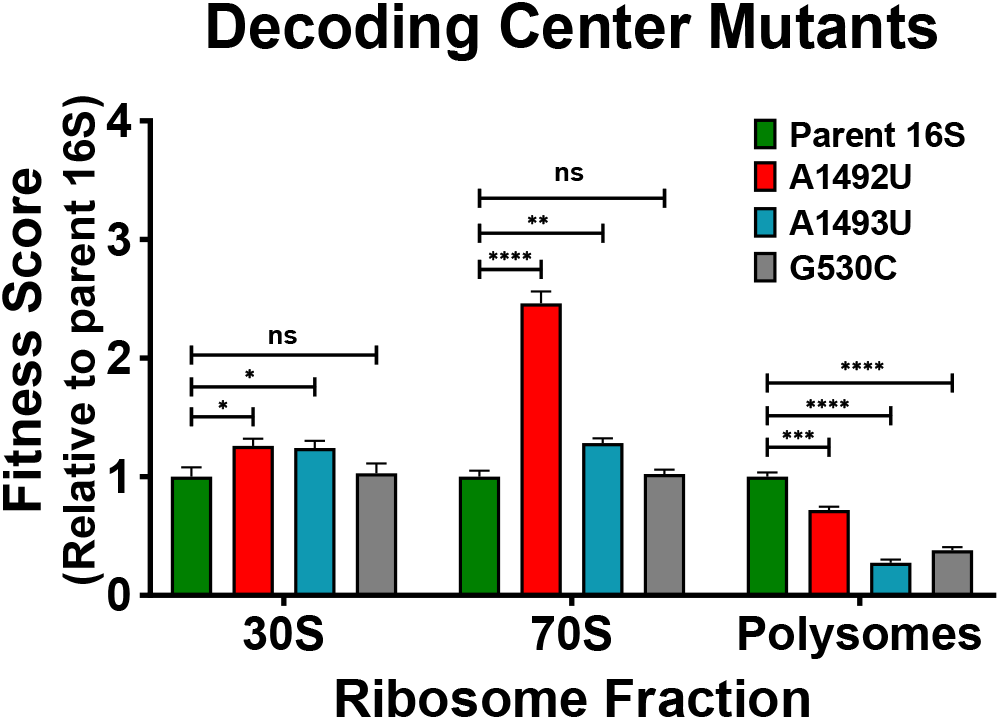
Performance assessment for decoding center mutants. Fitness scores were determined for plasmid-born 16S rRNA with separate decoding center mutations A1492U (red), A1493U (teal), and G530C (grey). Scores for mutants were compared to that observed for the parent 16S (y-axis) for the 30S, 70S, and polysome lysate fractions. Error bars represent standard deviations for biological replicates (n=3). Comparative statistics based on student’s *t*-test. *P* values <0.05 (*), <0.01 (**), < 0.001 (***), <0.0001 (****). *P* value ≥ 0.05 (ns).

### 4. Performance evaluation of disparate V3 region mutations

16S hypervariable regions from different bacteria tend to have similar sizes that symmetrically cluster around a mean value (34). One curious exception to this pattern is found in the V3 region encoding helix 17 (H17), the lower stem of which is involved in the assembly of the small subunit (35–37). The central region of V3 is highly variable and some bacteria have V3 versions lacking approximately 25 residues from the central portion, including members within a genus, *e.g., Clostridioides* (**Fig. 4A**) (34). This observation suggests that the central portion of V3 (encoding the outer stem-loop of H17) is not important for ribosome activity. However, we discovered that the central region of V3 is highly conserved in the class Gammaprotobacteria (which includes *E. coli)* (**Fig. 4B**). Moreover, individual 16S alleles within comparative reference strains have identical V3 regions (*E. coli* MG1655 *vs*. *Clostridioides difficile* 630), suggesting they are not entirely free to drift (**Fig. S3**). In the mature *E. coli* ribosome, the tip of H17 interacts with residues of the V2 region (**Fig. 4C**). Unfortunately, there is no available structure of a *C. difficile* ribosome for comparison. These striking contrasts in the architecture and conservation of the V3 region raised the question of its importance in ribosome activity.

**FIG 4:**
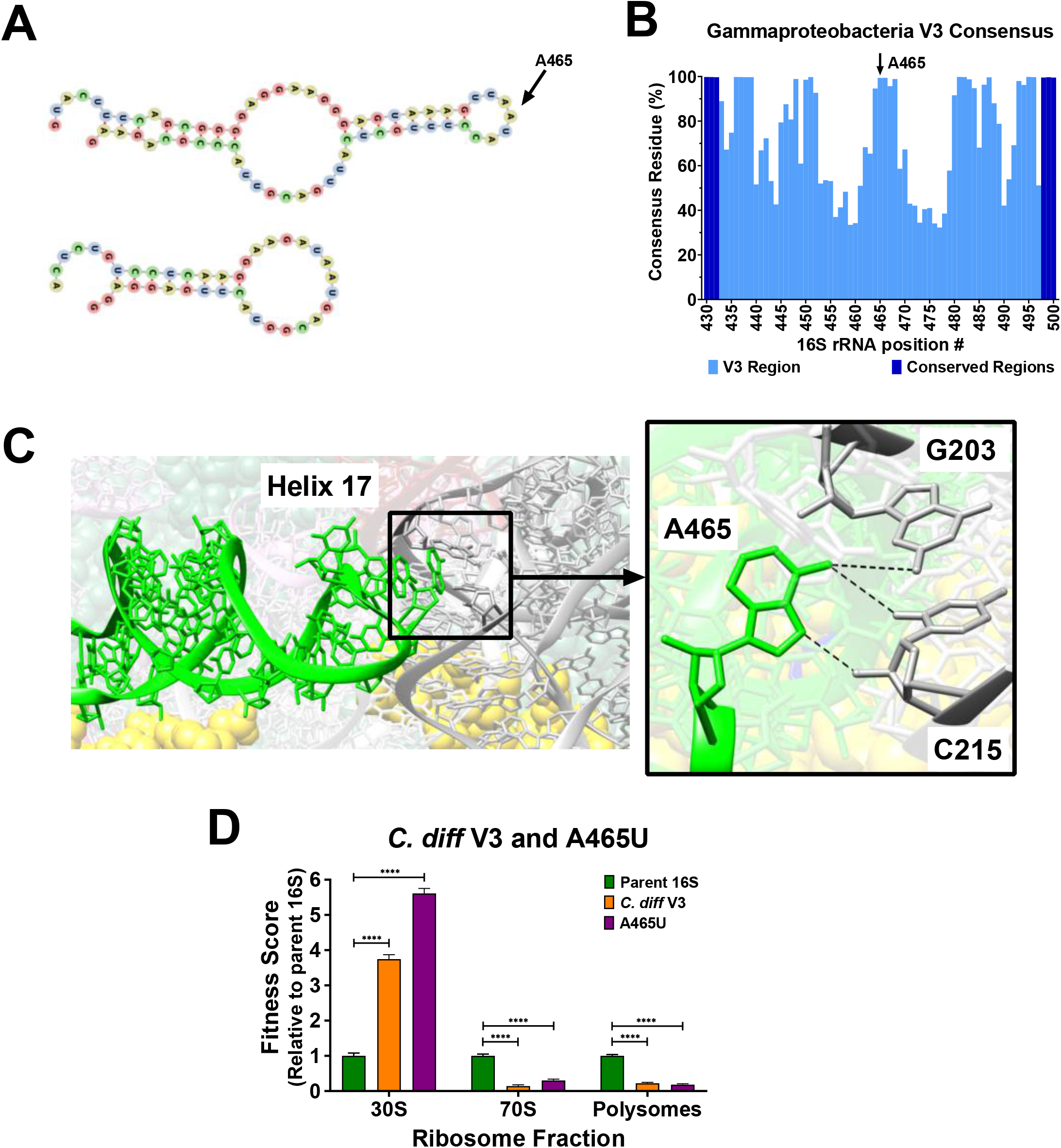
Identification and performance assessment of disparate V3 region variants. The central portions of V3 regions are generally not conserved and fall into two length categories. **A)** *Escherichia coli* and *Clostridioides difficile* V3 region secondary structures were computationally predicted using RNAfold (60). The *C. difficile* V3 encodes a shorter helix and is missing the outer stem-loop. At the tip of the *E. coli* hairpin is residue A465 (arrow). **B)** An analysis of residue consensus in V3 region sequences revealed that A465 is present in over 99% of organisms in the class Gammaproteobacteria (total of 162,325 sequences). **C)** In the *E. coli* ribosome, A465 is located at the tip of helix 17 (lime-green) and forms potential hydrogen bonds with G203 and C215 in the V2 region (dark grey). Yellow spheres are residues of small subunit proteins (image rendered from PDB 4V9D). **D)** Fitness scores for *E. coli* 16S rRNA with *C. diff* V3 (orange) and an A465U transversion (violet) were determined relative to the parent 16S. Error bars represent standard deviations for biological replicates (n=3). Comparative statistics based on student’s *t*-test. *P* value < 0.0001(****).

As an extreme evaluation of the importance of V3, we tested the performance of an *E. coli* 16S rRNA harboring the highly disparate *C. difficile* V3 region. The fitness scores for this variant indicated that most of the RNAs were trapped during subunit assembly, with very little progressing into the 70S or polysome pools (**Fig. 4D**). To interrogate the conserved V3 Gammaproteobacteria region, we evaluated a 16S version harboring a single SNP (A465U) that was predicted to alter the interaction between the H17 tip and the V2 region in the *E. coli* structure (**Fig. 4C**). Remarkably, this 16S variant performed nearly as poorly as the version containing the *C. difficile* V3 and much of it was retained in the 30S fraction (**Fig. 4D**). These findings support the hypothesis that conservation in the central portions of some V3 regions is due to a role in small subunit assembly that cannot be easily evolved. These data also highlight the potential for improving taxonomic classifications by focusing on the identities of individual residues that are hallmarks of particular groups of bacteria.

### 5. Positional relative entropy reveals strain- and species-specific residue variations

To evaluate the capability of SNPs to provide taxonomic information at a higher level than overall homology, we focused on interrogating the V3-V4 region because of its prominent use and extensive reference data sets from environmental and clinical studies. The *Escherichia* and *Shigella* V3-V4 regions typically have 1-4 residue differences among them, so they are of no utility during classifications based on homology. Prior studies evaluated the Shannon entropy associated with a consensus sequence across bacteria (10, 11, 38), but this method applied to strains or species would not weight SNPs that occur among alleles in a genome relative to the population. *Escherichia* and *Shigella* strains typically have seven copies of the 16S rRNA gene, each of which may have varying residues. To gauge the evolutionary change observed in a strain relative to the population, we used relative entropy (*D*_KL_) (39) to compare the residue frequency observed at a V3-V4 position across alleles within a strain to the frequency at that position for the overall population (**Fig. 5A**). Therefore, a residue variation in a single allele of only one organism of the population indicates less evolution, would yield a low *D*_KL_ value, and is less strain-informative. The same residue variation in all seven alleles of only one strain in the population indicates greater evolution, would yield a high *D*_KL_ value, and is more strain-informative. By taking the cumulative sum of *D*_KL_ (c*D*_KL_) at a position across strains in a population, we determined if the variation was prevalent in many strains and potentially species-informative.

**FIG 5:**
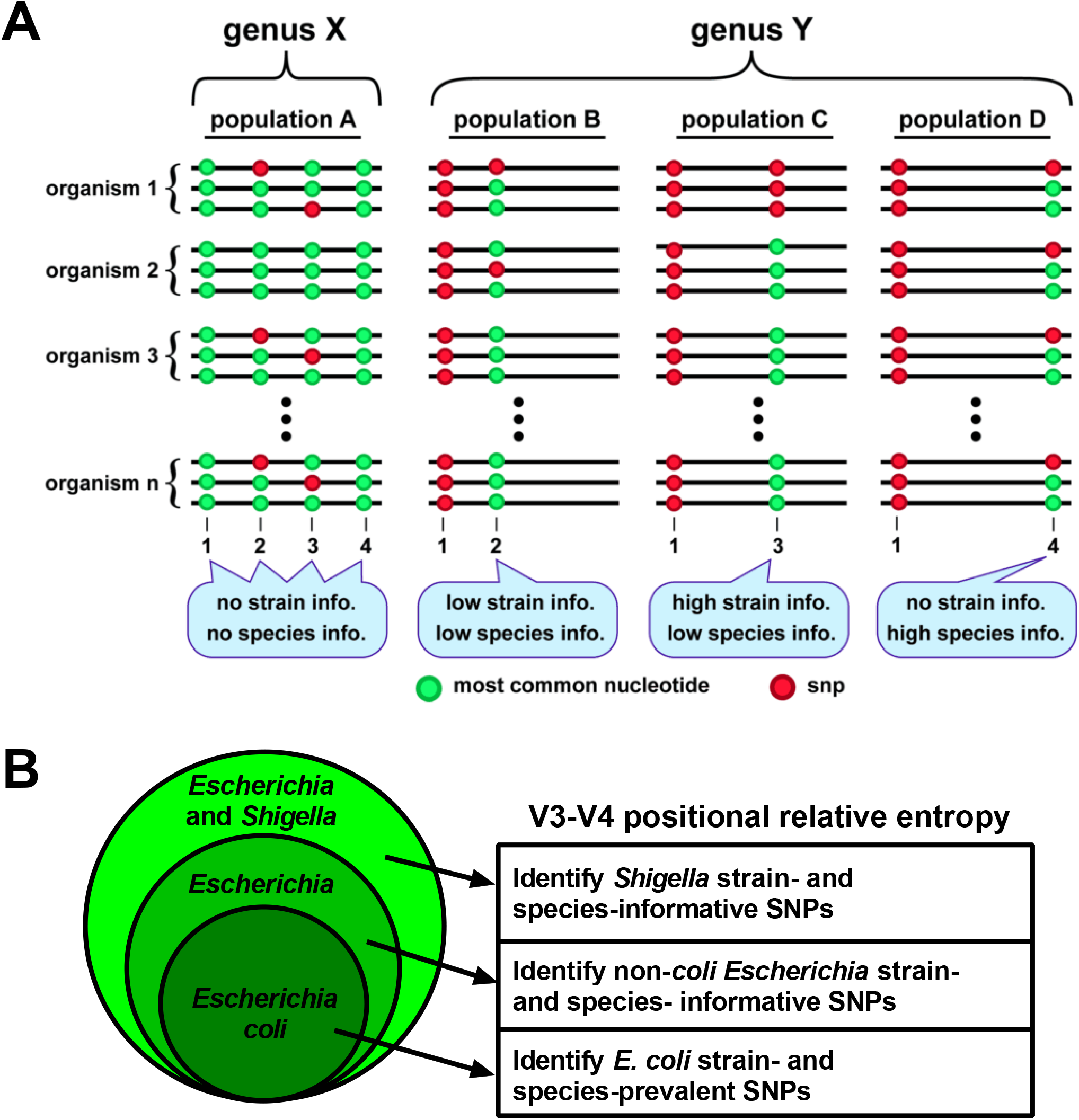
Illustration of V3-V4 positional relative entropy. Relative entropy analysis can be used to identify strain- or species-specific residues among variable region sequences in a population. **A)** Schematic depictions of variable regions of a multi-copy gene found in two hypothetical genera, X and Y. In a sequenced cohort of genus X organisms (population A), invariant residues at positions 1 and 4 provide no sub-genus information and SNPs observed at positions 2 and 3 are not associated with a particular species or strain, so their identities provide no information at those taxonomic levels. In genus Y, the SNP at position 1 (relative to that of genus X) is a strong genus indicator. In population B, occasional SNPs at position 2 provide no information because they are observed in single alleles in strains non-specific to a species. In population C, a SNP at position 3 is a strong strain indicator because it is present in all alleles in their genomes. In population D, occasional SNPs at position 4 indicate the presence of that species but provide no strain information. **B)** Positional relative entropy (*D*_KL_) of V3-V4 residues was employed to identify SNPs that were i) prevalent in a strain or multiple strains relative to the total *E. coli* population, ii) informative of *non-coli Escherichia* strains and species relative to the total *Escherichia* population, and iii) informative of *Shigella* strains and species relative to the total *Escherichia* and *Shigella* population.

We obtained V3-V4 region sequences from genomes of strains in the species *Escherichia coli,* the genus *Escherichia,* and the collective genera *Escherichia* and *Shigella*. We then computed the positional relative entropy of i) strains within the species *E. coli,* ii) *non-coli* strains within the genus *Escherichia,* and iii) *Shigella* strains within the total population of *Escherichia* and *Shigella* (**Fig. 5B**). Our use of relative entropy therefore identified residue variations that discriminated certain species within the closely-related genera *Escherichia* and *Shigella*.

### 6. Strain and species informative V3-V4 variants in *Escherichia* and *Shigella*

We employed positional relative entropy (*D*_KL_) to determine V3-V4 residues within *Escherichia* and *Shigella* that were strain- or species-informative. *D_KL_* values were determined comparing the observed residue frequencies at each V3-V4 position per *E. coli* strain relative to frequencies at that position across 12,876 V3-V4 sequences from 1,850 *Escherichia coli* strains. We plotted maximum *D*_KL_ values as well as the c*D*_KL_ values per position, which aids in identifying strain- and species-informative variations, respectively. As such, two SNPs were prominent across multiple *E. coli* strains (**Fig. 6A**), namely G474A and an absence of G666 (designated G666-). SNP G474A exhibited a *D*_KL_ of 4.373 and c*D*_KL_ of 296.478 and SNP G666-exhibited a *D*_KL_ of 4.01 and c*D*_KL_ of 163.856 which, when compared to theoretical values (**Fig. 6B** and **6C**), are prevalent in several strains and strongly informative of *E. coli* species.

**FIG 6:**
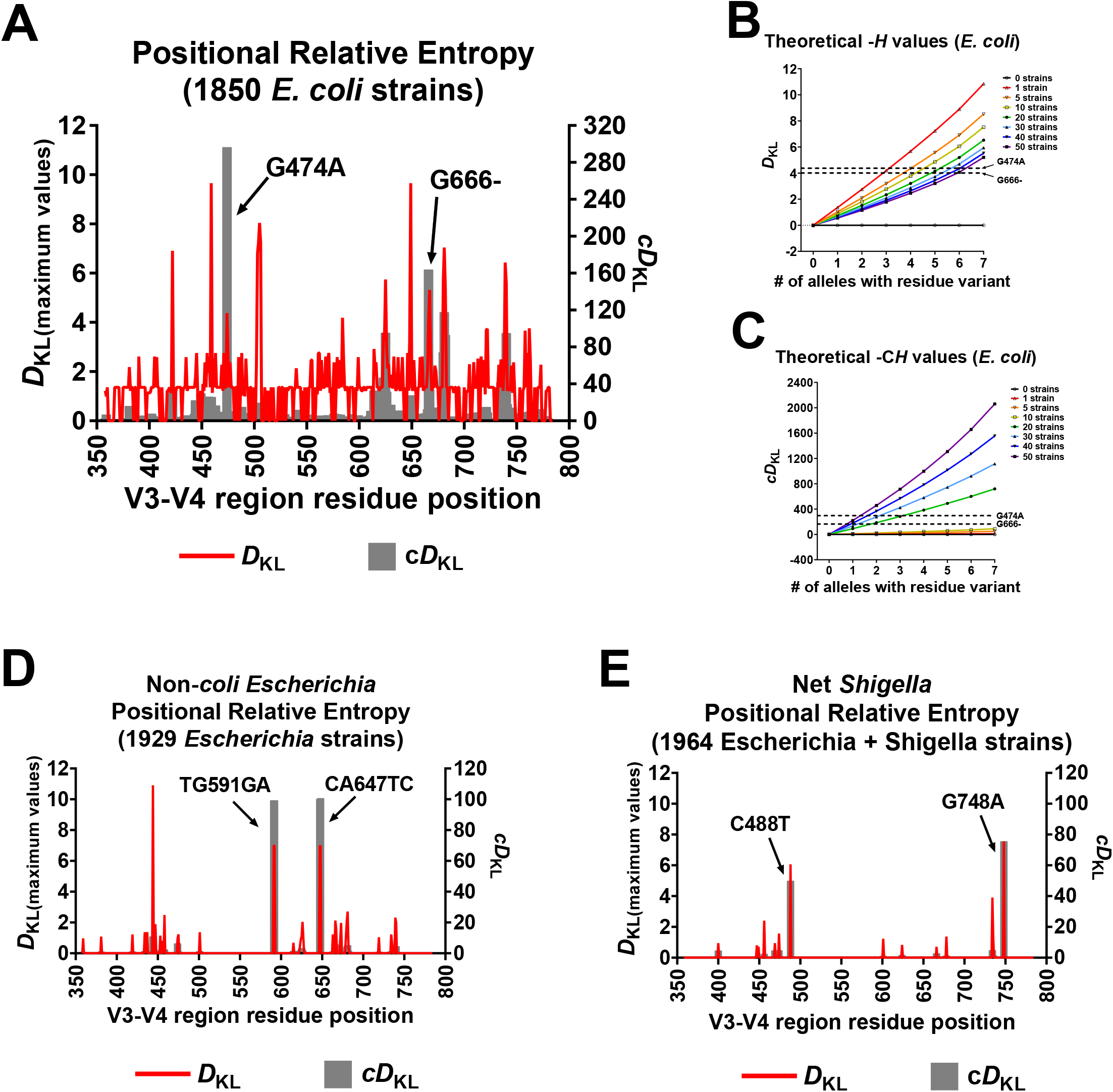
Informative V3-V4 sequence polymorphisms among *Escherichia* and *Shigella*. Relative entropy was used to identify strain- and species-informative SNPs within *Escherichia* and *Shigella* 16S V3-V4 sequences. **A)** *D*_KL_ peaks (red) correspond to SNPs at that position that are highly correlated with a particular strain among *E. coli* strains. Peaks in a cumulative *D*_KL_ (c*D*_KL_) plot (grey) indicate a SNP that is prevalent across strains and may be specific to *E. coli* species. For the evaluated *E. coli* population (1850 strains), two SNPs showed high species c*D*_KL_, namely G474A and G666. **B and C)** Theoretical values for *D*_KL_ and c*D*_KL_ were calculated for up to 50 strains in the *E. coli* population having a SNP in 0-7 out of 7 alleles. The *D*_KL_ and c*D*_KL_ values of the notable SNPs discussed in panel A are indicated for reference. **D)** *D*_KL_ values were determined for non-*coli Escherichia* strains. Two polymorphisms, TG591GA and CA647TC, had high strain and species values and were found only in *E. albertii* strains. **E)** *D*_KL_ was calculated to identify *Shigella* strain- and species-SNPs within the large population of *Escherichia* and *Shigella*. C488T and G748A had high strain and species values and were *S. boydii*- and *S. dysenteriae*-specific, respectively.

Relative entropy was also determined for V3-V4 positions per *non-coli Escherichia* strains considering the reference as positional residue frequencies across all *Escherichia* strains. (13,429 V3-V4 sequences from 1,929 *Escherichia* strains). Two sets of high *D*_KL_ and c*D*_KL_ values were observed corresponding to TG591GA and CA647TC (**Fig. 6D**), suggesting they are indicative of a strain and prevalent across strains. These polymorphisms are found in *Escherichia albertii* strains (total of 20) in multiple alleles in their genomes and are therefore strongly indicative of this species in a population of *Escherichia*.

Lastly, *D*_KL_ was determined for *Shigella* strains, considering the positional frequencies of residues in the total population of *Escherichia* and *Shigella* as references (1,964 strains). This analysis revealed two prominent SNPs with high *D_KL_* and c*D*_KL_, C488T and G748A (**Fig. 6E**). C488T is in most gene copies among *Shigella boydii* strains (total of 11) but was also in one or two gene copies among a few *E. coli* strains. G748A was found only in *Shigella dysenteriae* strains (total of 10) and was always present in all 7 copies of the 16S rRNA gene. Also, low *D*_KL_ values for G474A and G666-in the non-*E. coli* population sets indicates that these mutations were primarily present in the larger *E. coli* population. All relative entropy estimations and FASTA files of the V3-V4 sequences are in **Data Set S1**. Overall, positional relative entropy revealed species-informative V3-V4 residues for *E. coli, E. albertii, S. boydii*, and *S. dysenteriae*.

### 7. Ribosome performance for prevalent *Escherichia* and *Shigella* V3-V4 allele variants

A prior study reported that there was a translation activity difference for *Shigella* 16S rRNA in *E. coli* (32). We evaluated fitness scores for 16S rRNA with *Escherichia* and *Shigella* species-informative V3-V4 variations including variants observed in *E. coli* (G474A and G666-), *E. albertii* (TG591GA and CA647TC), *S. boydii* (C488T), and *S. dysenteriae* (G748A). Changes in fitness scores relative to the parent 16S illustrate that particular variable region changes can cause phenotypic effects.

*E. coli* ribosome structural studies showed that G474 is bonded with U458 (**Fig. 7A**) and G666 is a leading G in a poly-G stretch bonded with U740 (**Fig. 7B**). G474A showed fitness scores higher than WT for all fractions, while G666- showed an increase in the 30S score, and a decrease in 70S and polysome scores. These results indicate that the common *E. coli* V3 and V4 variants in 16S rRNA can perturb small subunit performance.

**FIG 7:**
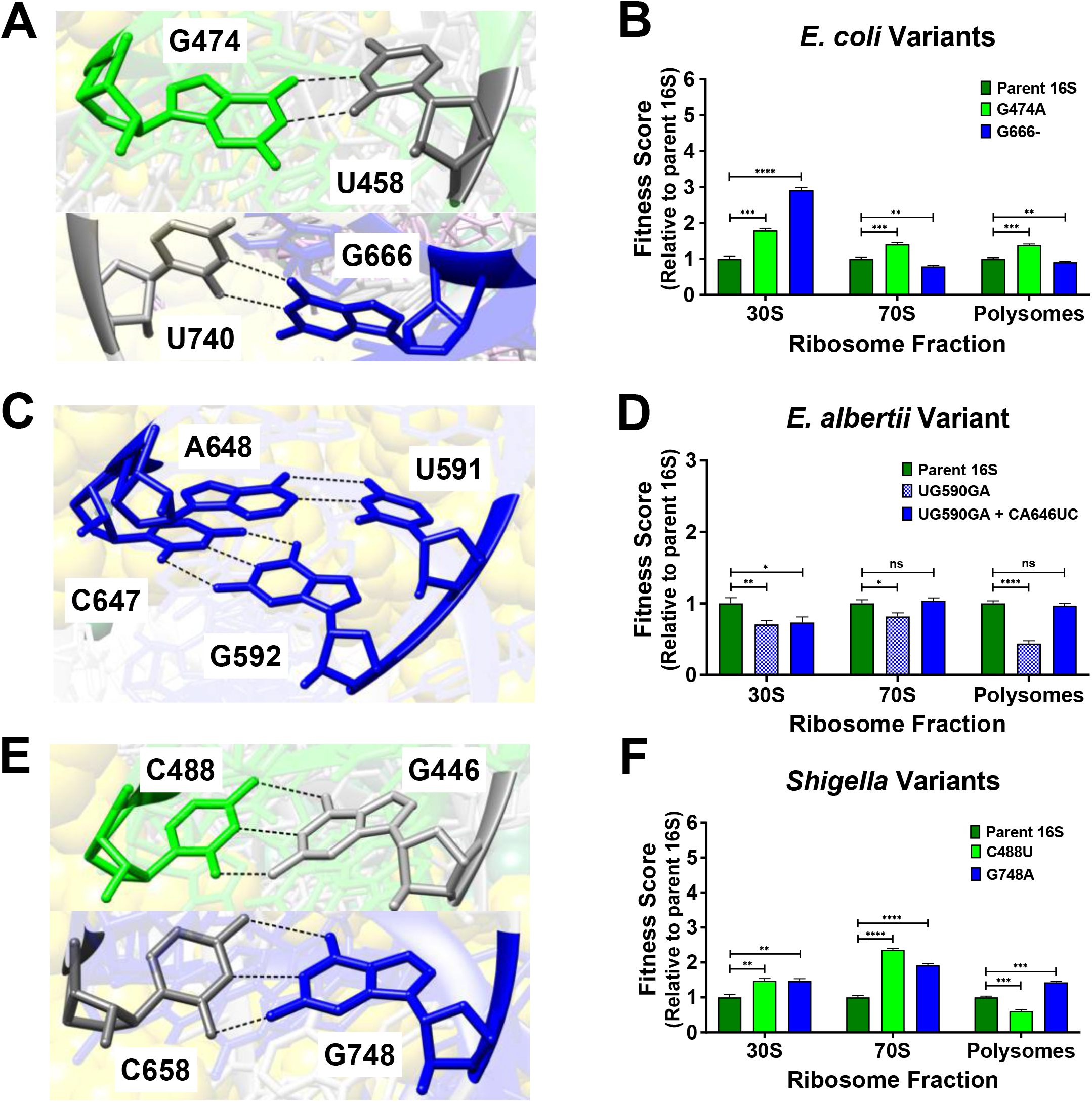
Structure of V3-V4 informative residues in *E. coli* and their fitness scores for *Escherichia* and *Shigella* species variants. The residue positions for *Escherichia* and *Shigella* species-informative SNPs were assessed in ribosome structures of *E. coli* str. K-12 substr. MG1655 (PDB 4V9D). Fitness scores were evaluated for *E. coli* 16S rRNA harboring informative *Escherichia* and *Shigella* species-informative V3-V4 residues. V3 residues in ribosome structures and fitness scores associated with their mutation were colored lime-green, and those for the V4 region were colored dark-blue. Residues that were not mutated were colored dark-grey in structures. **A)** G474 hydrogen bonds with U458 and G666 with U740. **B)** The fitness scores for 16S rRNAs harboring the species-informative *Escherichia coli* variations (G474A and G666-) were evaluated *in vivo*. **C)** Residues at sites for *E. albertii-specific* polymorphisms (UG591GA and CA647UC) complemented each other. **D)** Fitness scores for *E. coli* 16S rRNAs harboring *E. albertii* V3-V4 variant at one or both sites were evaluated. **E)** C488 and G748 are positioned to interact with G446 and C658 respectively. **F)** Fitness scores for *E. coli* 16S rRNAs harboring C488U (*Shigella boydii* SNP) and G748A (*Shigella dysenteriae* SNP) revealed that abundances in each were affected. Error bars represent standard deviations for biological replicates (n=3). Comparative statistics based on student’s *t*-test. *P* values <0.05 (*), <0.01 (**), < 0.001 (***), <0.0001 (****). *P* value ≥ 0.05 (ns).

*E. albertii* V3-V4 polymorphisms occurred at two sites tandemly (TG591GA and CA647TC). Ribosome structure studies showed that residues at these two sites form complementary hydrogen bonds (**Fig. 7C**). Observably, these *E. albertii* variations also complement one another. We evaluated fitness with a single site change (UG591GA) as well as both sites changed (UG591GA and CA647UC). The single-site change exhibited lower scores in 70S and polysome fractions, while the two-site mutation showed same scores as the parent 16S (**Fig. 7D**). Altogether, these results indicate that 16S rRNA abundances in translating ribosomes can be maintained so long as these residues co-vary. Because four nucleotide changes are required to convert an *E. coli* V3-V4 into this *E. albertii* version, and because an intermediate in this conversion is detrimental, there is a larger evolutionary hurdle to cross.

The *S. boydii* (C488U) and *S. dysenteriae* (G748A) polymorphisms would disrupt G-C Watson-Crick bonds found in the *E. coli* ribosome (**Fig. 7E**). Interestingly, G748A exhibited increased scores in polysome fractions (**Fig. 7F**), perhaps indicating more efficient translation initiation or slower egress from the translation pools. In contrast, C488U exhibited lower scores in polysome fractions, indicating that this variant was less capable to enter translation. Importantly, these results show that these *Shigella* species-informative V3-V4 SNPs have a biological impact on the performance of *E. coli* ribosomes, which indicates functional barriers to their evolutionary drift.

## Discussion

We characterized the performance of 16S rRNA variants with an eye toward establishing whether hypervariable regions contain biologically important residues. Interestingly, during the evaluation of our test system, we found that 16S rRNA with decoding center mutations were present in polysome fractions, suggesting they were capable of stably engaging mRNAs being translated by other ribosomes. This finding contrasts a previous study wherein similar 16S rRNA mutants were undetectable in the translation pool when overexpressed, indicating that perhaps subunit maturation had been compromised (29). Structural studies have elucidated the specific interactions of decoding center residues as they evaluate cognate tRNA binding (7, 40), but faulty interactions may not preclude ribosomes from progressing through translation initiation (29). Furthermore, the molecular interactions of these decoding residue mutants have not been established. Therefore, it is possible that small subunits with defective decoding centers can participate in translation, perhaps aberrantly decoding mRNAs. Further studies will be required to evaluate this idea.

We discovered that certain variable region 3 mutations altered the distribution of ribosomes in a manner suggesting they were unable to mature and escape the 30S pool. In the *Escherichia* and *Shigella* V3 sequences we computationally evaluated, an A465G SNP is present in a single *rrs* allele of only one *E. coli* genome and therefore is neither strain- nor species-informative. However, A465 variations, including U465, were found in 625 other Gammaproteobacteria V3 sequences (out of a total of 162,325), none of which belonged to *Escherichia* or *Shigella,* indicating that a residue other than “A” at position 465 is indicative of an organism divergent from these two genera. Although it was beyond the scope of this study, similar analyses can be conducted to evaluate other highly conserved residues among Gammaproteobacteria or other populations to uncover highly informative residues.

In contrast to A465, we identified individual V3-V4 residues that are informative of *non-coli* species within the genera *Escherichia* and *Shigella,* namely *Escherichia coli, Escherichia albertii, Shigella boydii,* and *Shigella dysenteriae. E. coli* 16S rRNAs harboring these species-informative variations exhibited fitness scores different from the parent 16S; however, when we introduced the *E. albertii* two-site covarying mutation, scores were restored to those of the parent 16S, which demonstrates the importance of co-variation between certain interacting regions to maintain biological function (26, 41). While we suggest that structural changes may be primarily responsible for changes in fitness scores, we note that covariation in 2D maps may not be sufficient to predict deleterious SNPs. For example, we observed that a 16S rRNA with a *Shigella dysenteriae* V4 residue exhibited higher fitness scores than the reference parent 16S even though a Watson-Crick bond was disrupted. This study was not designed to identify molecular mechanisms behind aberrant rRNA distributions, but it may be informative in future work to establish residence times of mutants within different ribosome pools to tease apart failures to mature (or to engage other translation factors) from changes in particle stability and lifetime.

Most bacteria have multiple copies of their rRNA operons (42), so a SNP that reduces ribosome performance may be tolerated if it does not interfere with the activity of the other ribosomes (for example, by stalling or titrating important factors). However, compensatory variations may occur in other parts of the ribosome which may explain, in part, why a strain can have multiple sequences among a hypervariable region, while a closely related strain has no variation among them. In our study, species-informative residue variations were found in the majority of 16S rRNA alleles in the genomes of respective species, indicating a greater evolutionary divergence compared to single allele changes. As such, we propose computational tools for establishing relatedness can be improved by incorporating a per-residue weighting metric that considers the multi-allele entropy derived from current sequence databases. It would still be necessary to confirm bacterial identity by other metrics (43–45), but computational re-evaluations of existing short-read sequence datasets may provide a higher confidence to the presence of particular organisms and offer more detail for evaluating community structures.

## Methods

### 1. Strains and plasmids

The *E. coli rrsA* locus was PCR amplified from strain BW30270 (Yale Stock Center, CGSC# 7925) initially using a forward primer that annealed in the upstream *hemG* gene to focus the amplification to that *rrs* allele. A second PCR was templated by the initial amplicon that generated a segment lacking the *rrsA* promoters, but retaining the native 5’ and 3’ processing regions, including the 3’ *ileT* and *alaT* genes (corresponding to residues 4,035,375 to 4,037,386 of the NCBI reference sequence NC_000913.3). This amplicon was then enzymatically assembled using NEBuilder (New England Biolabs, Ipswich, MA) with a DNA plasmid fragment such that *rrsA* expression was under the control of a *P_ara_BAD* promoter and a downstream *rrnB* terminator pair derived from plasmid pBAD24N (46). The complete p16S also contained a p15A replication origin and a *bla* selection gene. Mutations were introduced into p16S using PCR-based site-directed mutagenesis (47). The experimental host strain was the BW30270 derivative SM1344 (48).

### 2. Ribosomal 16S rRNA extraction and quantification

Experimental cultures were grown in MOPS EZ Rich Defined Medium (Teknova, Hollister, CA) (49) such that p16S was fully repressed, partially repressed, or fully induced. Triplicate cultures were harvested during exponential phase and cells pelleted at 4 °C. Cells were lysed by mechanical shearing in a cold lysis buffer that aided in keeping ribosomes bound to mRNA. Absorbances (at 258 nm) of replicate lysates were normalized. Normalized lysates were then separated across 10–40% sucrose gradients and fractionated while absorbance was measured at 258 nm. RNA was extracted from each gradient fraction of interest as well as from the total lysate using phenol/chloroform, which included a DNase treatment step. Complementary DNA was then generated for all RNA in extract samples. qPCR was then carried out using cDNA as templates and primers for tagged V1 or WT V1. qPCR fluorescence data were used to calculate plasmid-born and chromosome-born 16S rRNA abundance ratios and fitness scores (details of procedures in **Text S1**).

### 3. 16S rRNA sequence retrieval and relative entropy analysis of V3-V4 region sequences

The NCBI Assembly database was used to download *E. coli, Escherichia,* and *Shigella* feature table text files (50). Advanced filters for feature tables included “Latest RefSeq”, “Complete genome”, and “RefSeq has annotation”. A custom python script was then used to extract 16S rRNA gene locations by using keywords “16S ribosomal RNA” and “ribosomal RNA-16S”. Thereafter, using a custom python script that incorporated the NCBI efetch tool, the “+” strand sequences of all 16S rRNA gene copies were obtained as a FASTA file. Multiple sequence alignments were carried out using MUSCLE (51). Aliview (52) was then used to extract the V3-V4 segments of the alignment, contained between the 341F and 785R primer binding sites (8, 53). Column gaps observed for all *E. coli* str. K-12 substr. MG1655 16S sequences across the alignments were removed (NCBI RefSeq# NC_000913.3). A python script incorporating Kullback–Leibler divergence (*D*_KL_) was then executed to determine the V3-V4 positional relative entropy for observing a residue across alleles in a strain compared to the entire population (**Equation 1**) (39, 54). The output file contained *D*_KL_ per residue position for each strain in the respective populations of study (**Table S1**).

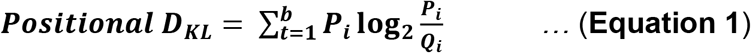

Here, *D*_KL_ values are position-specific in the sequence alignment. *P_i_* is the frequency of the observed *t*^th^ residue (A/C/G/T) at the *i*^th^ V3-V4 position across alleles in a strain. *Q_i_* is the reference frequency of that residue at that V3-V4 position across all sequences in the population of study. *D*_KL_ was used to identify variations specific for *E. coli* strains in the population of *E. coli, non-coli Escherichia* strains in the population of *Escherichia*, and *Shigella* strains in the population of *Escherichia* with *Shigella*. For nucleotides with 0 occurrence, 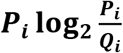 was evaluated as 0. A computed value of 0 implies that there is no positional difference in the residue between the strain and the overall population. A high *D*_KL_ value is indicative of a different residue in multiple alleles of a strain at that position relative to the majority of the population. A cumulative sum of *D*_KL_ (c*D*_KL_) at a position determined if that variation occurred across multiple strains. Designations for strains that contributed to high c*D*_KL_ values were determined if they were a common species.

### 4. V3 sequence determination of Gammaproteobacteria and *Clostridioides difficile*

A premade multiple sequence alignment of non-redundant Gammaproteobacteria 16S rRNA sequences was retrieved from SILVA Ref NR 99 Release 138.1 (55). Jalview (56) was used to obtain the consensus residue percent for the V3 region (8) between residue 433-497 using *Escherichia coli* MG1655 as a reference. Separately, 16S gene sequences from *Clostridioides difficile* str. 630 (NCBI RefSeq# NZ_CP010905.2) and *Escherichia coli* str. K-12 substr. MG1655 (NCBI RefSeq# NC_000913.3) were obtained and aligned using Aliview. Alignment positions corresponding to residues 433-497 for the *E. coli* reference 16S gene were considered as the *C. difficile* V3 region.

### 5. Structure analysis

Chimera X was used to view and evaluate *E. coli* ribosomes (57). All illustrations used PDB 4V9D which contains a ribosome with bound tRNA and is commonly used to determine RNA-RNA interactions in *E. coli* 16S rRNA (58, 59). Hydrogen bonds were illustrated using default parameters, ignoring intra-residue bonds. Putative V1 and V3 region secondary structures were determined using RNAfold (60). The predicted tagged V1 secondary structure was similar to a biochemically established *E. coli* V1 secondary structure (59).

## Abbreviations

SNP: Single Nucleotide Polymorphism
RT-qPCR: Reverse Transcription and Quantitative Polymerase Chain Reaction
*D*_KL_: Kullback–Leibler divergence

## Acknowledgement

We thank Dr. Shibu Yooseph, Dr. Taj Azarian, and Michael Johnstone for their editorial comments. This work was supported by NIH Grant 1R01GM118896.

---

**FIG S1: V1 region tagging strategy for qPCR and specificity of qPCR primers.** The V1 region of *E. coli rrsA* in our p16S was modified with a tagged sequence. **A)** Secondary structure the tagged V1 vs. WT V1 for 16S rRNA were found to be similar (compared using RNAfold (60) and modeled towards *E. coli* 16S rRNA V1 region helix 6 (59)). qPCR primers were designed to bind either to the tagged V1 or WT V1 (primer binding sites illustrated as solid red and solid green lines, respectively). Sequences were added to 5’ tails of qPCR primers (black and orange zigzags) and a primer binding site added in the forward primer for Sanger sequencing (blue line). The total qPCR product length was expected to be 120 bp (see table S2 for primer sequences). **B)** The specificity of qPCR primers to detect purified plasmid tagged V1 *rrsA* and WT V1 *rrsA* was determined. ΔCq for amplification of specific vs. non-specific target was plotted (x-axis) for each template (y-axis). Error bars represent standard deviation for biological replicates (n=3).

**FIG S2: Growth dynamics of WT cells with controls.** Growth dynamics for *E. coli* cultures with parent p16S and plasmid controls were assessed when expression was uninduced and induced. Plasmid controls tested include p16S, but with i) the 16S gene removed (Δ16S), ii) *rrsA* with untagged V1, and iii) *rrsA* replaced by a *gpH* ORF fragment the same size as the 16S rRNA gene. Log2 values of absorbance at 600nm (y-axis) is plotted against time (x-axis). Log2 values were shifted relative to uninduced p16S to aid visualization. Growth rates for all were estimated between 435 and 455 min, during late exponential phase. Error bars represent standard deviation for biological replicates (n=3). Comparative statistics based on student’s *t*-test. *P* value < to 0.001 (***).

**FIG S3: *E. coli* and *C. diff* V3 region alignment**. Aliview was used to perform multiple sequence alignment of 16S gene sequences from *Clostridioides difficile* str. 630 (NCBI# NZ_CP010905.2) and *Escherichia coli* str. K-12 substr. MG1655 (NCBI# NC_000913.3). The V3 region is highlighted in a box. Residues are colored as: A in green, C in blue, G in black, and T in red. Alleles were identical among genomic copies for respective strains and so a pairwise alignment of a representative sequence from each strain was used to perform a pairwise alignment using the Needleman-Wunsch algorithm.

**Table S1: qPCR primer sequences targeting tagged V1 and WT V1 16S sequences.**

**Data Set S1: Strain list, V3-V4 sequences from species *E. coli,* genus *Escherichia,* and collective genera *Escherichia* and *Shigella,* and relative entropy results.**

**Text S1: Methods detailing culturing harvesting and lysis, lysate fractionation, RNA extraction, and RT-qPCR.**

